# A microfluidic platform to investigate the role of mechanical constraints on tissue reorganization

**DOI:** 10.1101/2022.03.16.484527

**Authors:** Sham Tlili, François Graner, Hélène Delanoë-Ayari

## Abstract

Mechanical constraints have a high impact on development processes, and there is a need for new tools to investigate the role of mechanosensitive pathways in tissue reorganization during development. We present here experiments where embryonic cell aggregates are aspired through constrictions in microfluidic channels, generating highly heterogeneous flows and high cell deformations that can be imaged using two-photon microscopy. This approach provides a way to measure *in situ* local viscoelastic properties of 3D tissues and connect them to intracellular and intercellular events such as cell shape changes and cell rearrangements. Perspectives include applications on organoids to investigate and quantify rheological properties of tissues, and to understand how constraints affect development.

## 1 Introduction

### 1.1 Motivation and state of the art

During embryo morphogenesis or wound healing, biological tissues are sculpted by active processes such as cell growth and division, cell migration or cell differentiation (1). These processes generate mechanical stresses within the tissue that can trigger tissue scale flows, cell deformation and rearrangements. The nature of the tissue mechanical response to mechanical stresses depends on its rheological properties at the scale of a cell group (or “cohort of cells” (2), or “representative volume element”). In turn, these cell group scale properties arise from the cell scale interplay between cells viscoelasticity and cell-cell interactions such as adhesion.

To fully understand and measure these properties, 3D *in vitro* tissues such as cancer cell spheroids or embryonic organoids (3) play an increasingly important role in biophysical and medical studies as they are easy to handle and their properties are relevant to better understand *in vivo* tissues.

To study this *in vitro* tissue rheology in the context of developmental biology, several experimental setups have already been used, including cell monolayers stretching (4), rounding of cell aggregates (5–7), shearing of cell monolayers or cell aggregates (8–11), and compression of cell aggregates (6, 12, 13). These experiments have enabled to measure bulk parameters such as shear and compression moduli, viscosity, and yield deformation or yield stress. To connect these mechanical properties to cellular processes, it is essential to have access at the same time to the cell structure and to its dynamics (e.g. cell shapes or rearrangements) (14).

Live imaging *in vivo* (during embryo development or insect metamorphosis, for instance) or *in vitro* gives the opportunity to track cell movements and divisions, cell shape variations and cell rearrangements, and to quantify their contributions to the cell group or tissue scale dynamics, by measuring their contributions to the cell group or tissue deformation rate for example (15, 16).

Methods combining the imposition of an external stress, force, deformation, or deformation rate controlled by the experimentalist with live imaging at a sub-cellular resolution have proven to be powerful approaches to investigate biological tissues and more generally complex materials rheological properties. For example, cell monolayers can be stretched by being suspended between rods or on an elastic membrane while imaging the tissue response at the cell level (17, 18). Nanometric magnetic beads naturally endocytosed by the cells have also been used to apply a global force on cell aggregates (13, 19).

A fruitful approach for 3D tissues is to embed ferro-magnetic droplets within the tissue which can be actively deformed by an externally applied magnetic field while imaging the tissue response at the cell scale. This method has enabled to measure spatial gradients of yield stress leading to a liquid-solid transition in the zebrafish mesoderm, and gives also access to the visco-elastic response of cells at short time scales (20). This approach provides a powerful method to probe locally tissue rheology at small deformation. To measure effective properties at large deformations, pipette aspiration are broadly used on 3D tissues such as cancer spheroids (10), embryos (e.g. mouse, Xenopus (21), chicken (22)) and explants. However, it is not easy to image the tissue microstructure and its dynamical response in such aspirations geometries. Cell-cell rearrangements are of high importance in developmental biology and are rarely observed in experiments in 3D. There was a need to design a system where they could be properly quantified so as to understand which biophysical principles govern their dynamics.

### 1.2 Approach and experiment principle

In this paper, we present a method to study 3D tissues cell group scale mechanical properties and to relate these properties with cell scale structure such as cell shape and heterogeneities in cell proteins and gene expression. This approach is inspired from studies on non-living amorphous materials (23, 24). Our method combines microfluidic aspirations with two-photon imaging of cell shapes at subcellular resolution. It enables to determine and correlate the spatiotemporal patterns of deformations generated within the 3D tissue, the visco-elasto-plastic properties of cell groups, and the dynamics of cell rearrangements.

The aspiration geometry, forcing a 3D cell aggregate to flow through a constriction, enables to apply a heterogeneous flow. As a consequence, each individual experiment provides a self-sufficient data set with a large enough variation range of cell group deformations and deformation rates. Moreover, this geometry enables to get large deformations and to change tissue boundary conditions to probe tissue elasto-capillary behaviour.

We pre-confine the aggregate and use a capillary with straight sides (rather than a cylindrical pipette). The tissue is physically constrained in the vertical dimension corresponding to the optical axis. This forces it to flow within the plane of observation (Fig. 1, Supp. Movie 1). Such quasi-2D flow facilitates imaging cell shapes and rearrangements.

**Figure 1:**
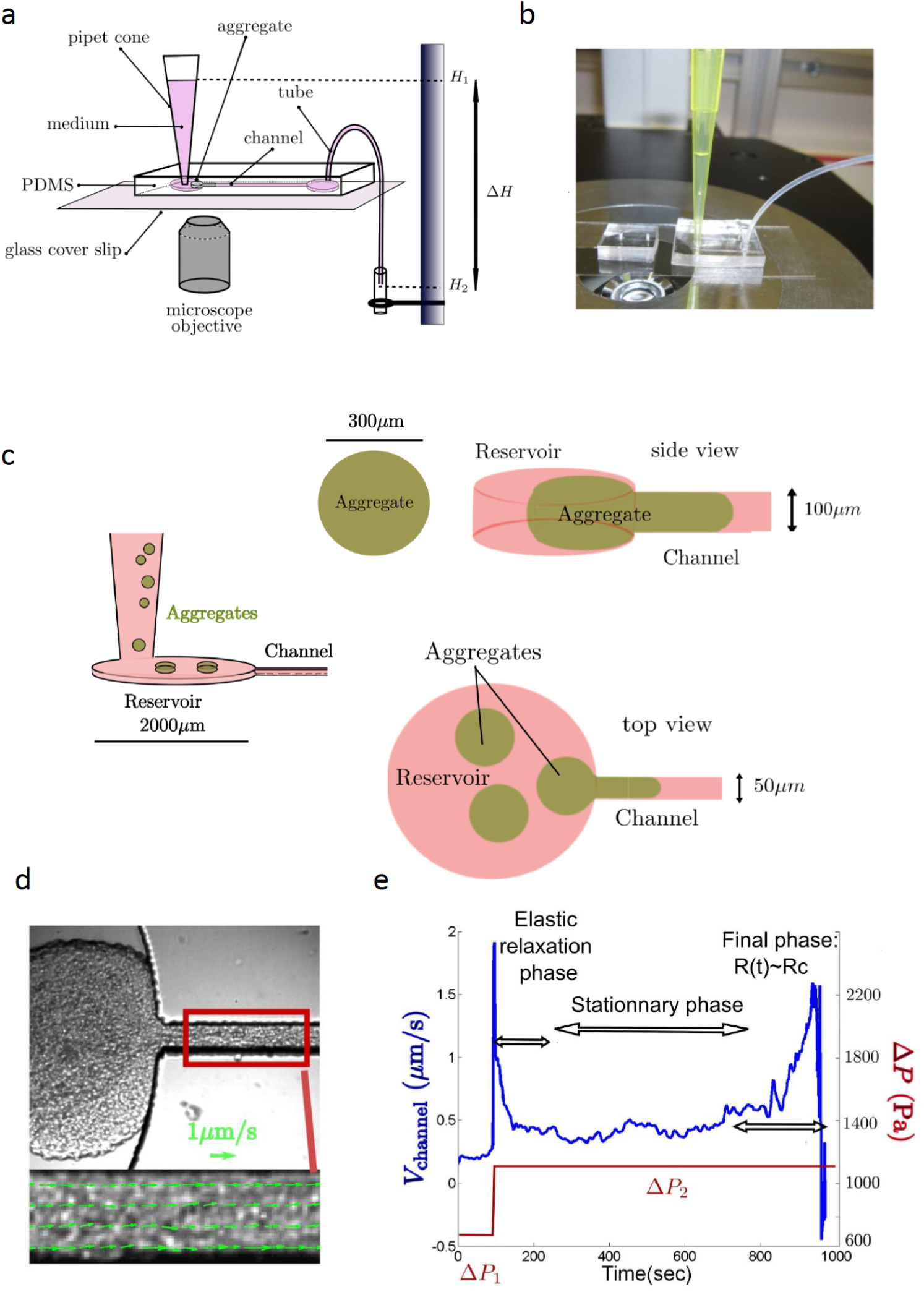
Aggregate aspiration. (a,b) Microfluidic device used to aspirate aggregates. Scheme and photograph of the device: a hydrostatic pressure difference is applied on the aggregate blocked inside the microfluidic channel. The microfluidic device is on a glass coverslip that allows imaging with an inverted microscope. (c) Microfluidic reservoir and channel top and side views. Preconfinement of the aggregate before aspiration: aggregates are put into the pipet cone, they sediment towards the channel reservoir. A pressure difference is applied inside the channel; the aggregate starts to enter into the reservoir and to deform in a pancake shape. (d) Brightfield image of an aggregate aspired in a channel (Supp. Movie 1). The red rectangle is the region used to quantify the flow rate. The inset shows the plug flow inside the channel which we could spatially average to obtain the mean velocity. (e) Aggregate flow rate inside the channel versus time.

In principle, the flow could have a 3D structure, namely depend on the direction *z* perpendicular to the device plane. This would be the case for instance for an aggregate with a core-shell structure, or any heterogeneous tissue. In that case our set-up would be suitable to perform a 3D image analysis, provided the flow was slow enough. In the following, we present experiments performed on homogeneous 3D aggregates of cells which do not have core-shell structures (Fig. 9, Supp. Movie 2), and we check here that we can neglect the flow variation along *z* (Supp. Movie 3).

## 2 Material and Methods

### 2.1 Manipulation

For these experiments, we used a mouse embryonic carcinoma F9 cell line (25). Aggregates were obtained by resuspending single cells after trypsinization in a dish and preventing them to settle down using an orbital shaker. After two days we obtained aggregates with a diameter around 250 *μ*m.

The experiment is performed on a microfluidic chip, Fig. 1a-c. The PDMS mold was obtained using standard lithography techniques. For the experiment presented here, the channel width was 50 *μ*m and its height 100 *μ*m. After bonding the PDMS chip to a glass coverslip using a plasma, we incubate the channel for 30 min with a Pluronic solution in order to avoid the adhesion of the aggregate to the wall while flowing. It is then rinsed with Leibovitz complete medium, which was further used for the experiment. For the introduction of the aggregate we use a 200 *μ*l cone, and the outlet is then linked to a fluid reservoir whose height can be adjusted in order to have a difference of pressure controlled by the difference in height between the sample and the reservoir.

The aggregate is first dropped into a cone pipette. Then it sediments into a 100 *μ*m high reservoir and deforms into a pancake shape (Fig. 1a-c). After waiting at least more than 10 minutes for stress relaxation, the aggregate is pushed towards the channel entrance. Because of the applied pressure difference, the aggregate slides into the reservoir towards the channel entrance and starts to deform inside the channel (Fig. 1d).

The aggregates were imaged on a two-photon inverted microscope using an oil immersion 40X objective (HCX PL APO CS 40.0×1.25 OIL UV) equipped with MaiTai Spectra Physics laser (690-1060 nm) used at 800 nm, and a lowpass filter below 600 nm, under a temperature control environment setup at 37^*°*^. To visualize the cell contours we used a solution of sulforhodamine B (6). We diluted 16 *μ*l of a stock solution (1 mg/ml) in 1 ml of Leibovitz complete medium. With this method, bleaching is very low as the fluorophore rapidly diffuses from the medium to the extracellular space.

### 2.2 Image analysis

We measure the two-dimensional velocity field 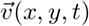 using a custom-made Matlab optic flow code based on the Kanade-Lucas-Tomasi (KLT) algorithm (26) with a level 2 pyramid. This coarse-grained velocity field can be averaged in time, yielding components *v_i_* (*x_j_*) where *i,j* = 1 or 2. Using finite differences we obtain in each box the deformation rate as the symmetric part of the velocity gradient, *grad* 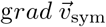, with components *(∂_i_v_j_* + *∂_j_v_i_*)/2. This symmetric tensor can be diagonalized, we graphically represent in each box the tensors by bars which direction and length correspond respectively to eigenvectors and eigenvalues, with blue bars for compression and red bars for elongation (Fig. 10).

To extract the coarse-grained cell anisotropy, we use Fourier Transform; details and validations are provided in (27). In short, the Fourier Transform of an image of a connective tissue estimates the cell size, shape and anisotropy averaged over a cell group. This method provides an efficient measurement of the deformation coarse-grained at cell group scale, without having to recognize and segment each individual cell contour. We use the same grid and box size as for the velocity (box of 256 pixels, overlap 0.75). To improve the signal-to-noise ratio, the Fourier Transform norm is smoothed in time by using a sliding average over three consecutive time-points.

We define the cell shape deformation tensor *ε*_cell_ with respect to a rest state which we assume to be isotropic, with two equal eigenvalues *L_0_* = (*L*_max_*L*_min_)^1/2^. Hence *ε*_cell_ has the same eigenvectors as the Fourier pattern, and two eigenvalues 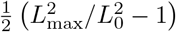 and 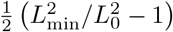. The deformation tensor *ε*_cell_ιι is equivalent to other definitions of the deformation (27, 28) within a linear approximation; in addition, *ε*_cell_ has the advantage to have well established transport equations (29).

We represent the deviator 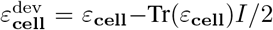, where *I* is the identity tensor; 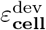 is the anisotropic part of *ε*_cell_ and only depends on the ratio 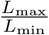, which is represented by ellipses whose color indicate the value of the deformation.

## 3 Results

### 3.1 Aspiration and brightfield microscopy

Here, we first focus on cell group scale aspects of the flow properties. Using brightfield or phase contrast microscopy gives already valuable information and enables to map the velocity field and deformation rate in the aspired aggregate. The velocity profile inside the channel is consistent with a plug-flow (Fig. 1d, inset) and we integrate it over the channel cross-section to measure the aggregate flow rate. Over time, we observe three distinct phases of the aspiration (Fig. 1e): (i) a rapid elastic aspiration phase which corresponds to the rapid entrance through elastic cell deformation of the aggregate inside the channel; (ii) a stationary phase where the flow rate is rather constant, corresponding to an effective viscous behavior of the tissue; and (iii) a final phase of acceleration when the non-aspired aggregate part size is similar to the channel size, which indicates that the dissipation on the channel walls is negligible compared to the viscous dissipation inside the aggregate.

Taking advantage of the pseudo-2D flow structure, we quantify the in-plane spatial variation of the flow and deformation rate within the aggregate (Fig. 2a). Models (10) predict that in a pipette experiment, the dissipation occurs only in a typical volume of effective radius R_channel_. In our setup, we can have a spatially resolved measurement of the deformation rate: in our geometry, the typical size of the dissipation zone is similar to R_channel_, but we observe an extended zone along the walls of the chamber. We then measure the velocity gradient, symmetrize it to determine the cell group deformation rate, and plot its anisotropic part (Fig. 2b). The isotropic part of the cell group deformation rate (Fig. 2c) shows a compression against the walls before the channel entrance and an expansion after the entrance of the channel, which illustrates tissue compressibility.

**Figure 2:**
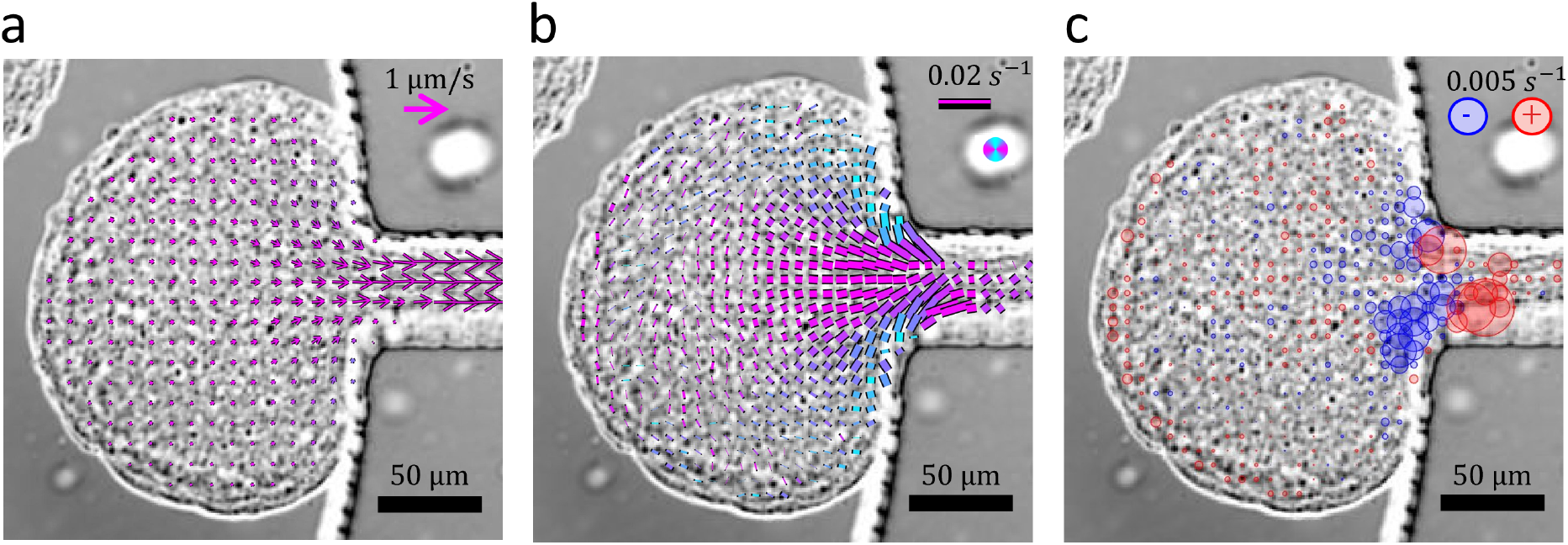
Aspiration: maps of velocity and deformation. (a) Time averaged velocity field over 5 min. (b) Anisotropic part of cell group deformation rate. (c) Isotropic part of cell group deformation rate. Scales are indicated and measurement maps are overlayed with brightfield images.

### 3.2 Cell deformations and rearrangements during aspiration

To investigate the link between the material properties and the cell dynamics, we image the tissue at the cell scale to measure cell deformation. Here we use the same formalism that we used previously on a 2D system (MDCK monolayers (30)). Cell divisions can be neglected when they are inhibited, or when the timescale of the experiment *T_exp_* is much smaller than the timescale of cell cycle T_div_; this is the case here, where T_exp_ ≈ 1 hour while T_div_ ≈ 12 hours. Then the cell group (total) deformation rate 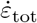 is the sum of the cell deformation rate coarse-grained at cell group scale, 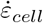, and plasticity deformation rate 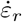 due to the rearrangement rate. The cell deformation pattern is due to the balance between: (i) the deformation induced by the constriction (i.e. the source term *grad* 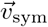); (ii) the relaxation of cell deformation due to cell rearrangements 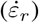; and (iii) elasticity transport terms (i.e. the fact that non-deformed cells are constantly advected towards the channel). We want to estimate the effective viscoelastic relaxation time related to cell rearrangements *τ_r_* (Fig. 3), which can be defined as 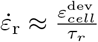 where the superscript dev indicates the deviatoric (i.e. anisotropic) part of a tensor.

**Figure 3:**
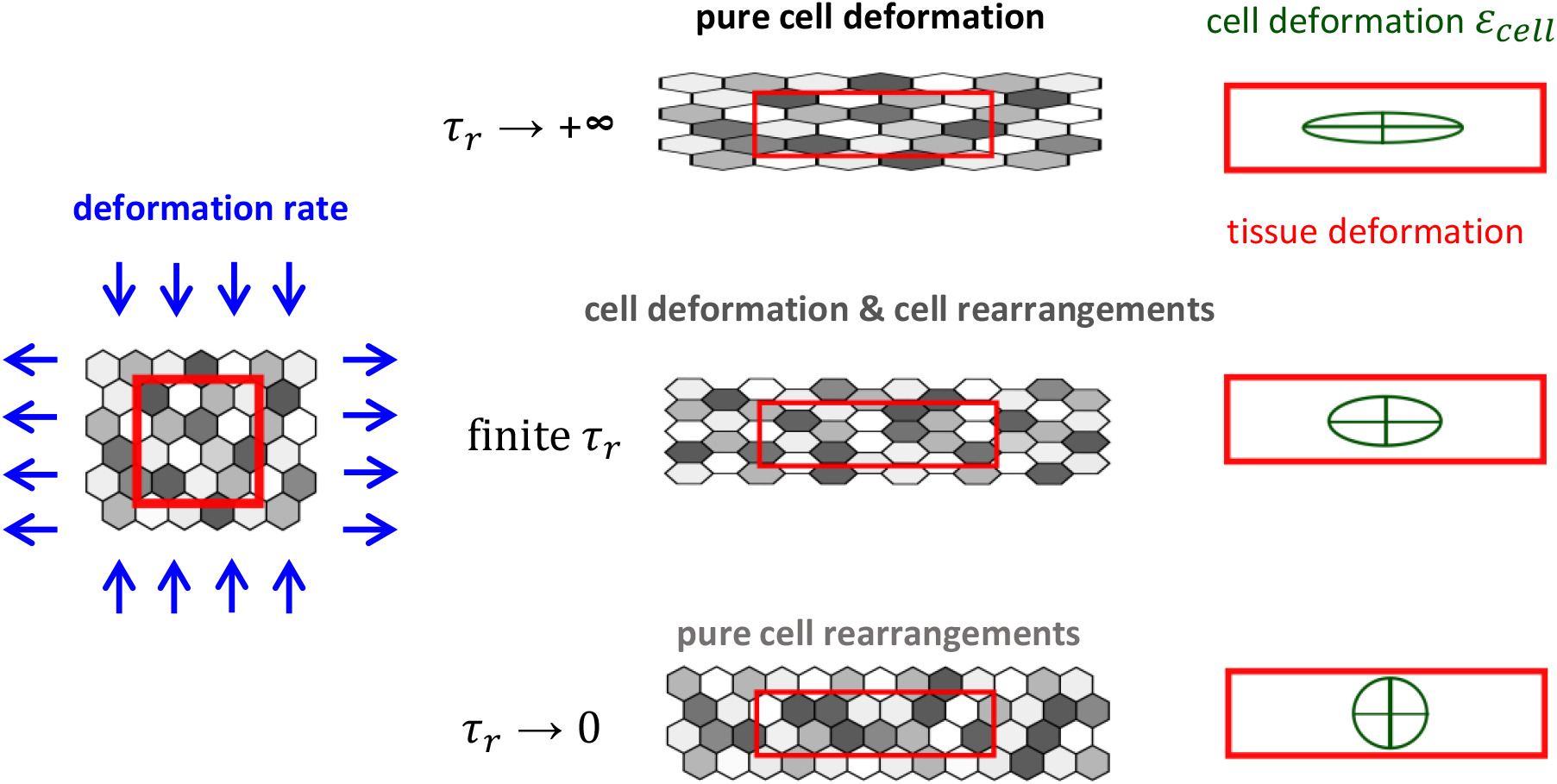
Effect of deformation rate on cell deformation. The deformation rate is shared between the elastic deformation rate, which results in cell deformation, and the plastic deformation rate, which results in cell rearrangements (15). Their respective shares are determined by the typical timescale *τ_r_* over which the deformation relaxes due to rearrangements. In the extreme limit of no rearrangements (top, *τ_r_* → ∞) cells only deform, and in the opposite limit of instantaneous rearrangements (bottom, *τ_r_* → 0) cells do not deform at all.

We image at sub-cellular resolution the process of aspiration and observe simultaneously the heterogeneous velocity and deformation fields (Fig. 4a, Supp. Movie 4). To take into account the transport of cell elasticity in the evolution of cell deformation, we measure the spatially averaged cell deformation 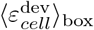 and the deformation rate 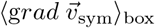 in a virtual rectangle advected and deformed by the tissue flow (Fig. 4b) using a home-made code. We can thus evaluate the terms of the evolution equation:

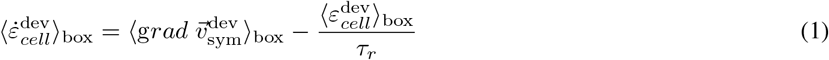

**Figure 4:**
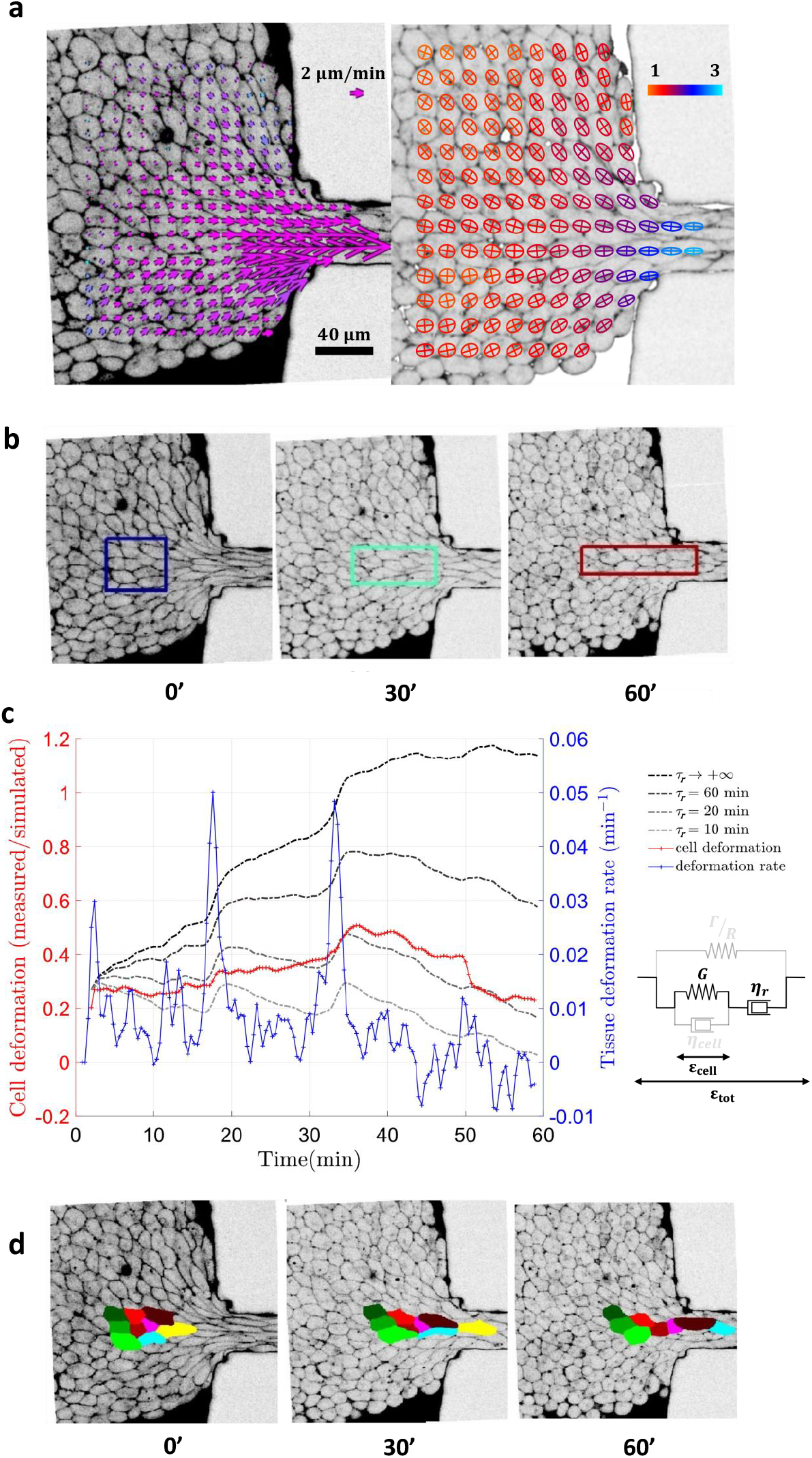
Quantifying cell shape changes and relaxation during the aspiration. The aggregate is imaged with an inverted two-photon microscope. (a) The optic flow method and Fourier transform method are performed on the cell contours (visible with the sulforhodamine B dye) to measure the velocity field (on the left, average over 5 minutes), and the ellipses representing cell group shape anisotropy (on the right, average over 5 minutes). Anisotropy is color-coded from 1 (red) to 3 (blue). (b) Virtual patch of tissue advected and deformed by the velocity field and its gradient, measured by optic flow (Lagrangian approach). This patch is used to spatially average the tensors in the following analysis. (c) Estimation of the relaxation time *τ_r_*. In blue, anisotropic component of cell group deformation rate 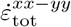, in red 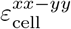, in gray levels simulated 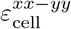 for different typical values of cell shape relaxation timescale *τ_r_*, in black simulated 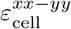 if there was no cell shape relaxation (purely elastic deformation, *τ_r_* → ∞). (d) Tracking of a cell group in the virtual patch and identification of cell rearrangements. Times are indicated in minutes.

We can measure at each timepoint the box-averaged tensors ⟨*ε_cell_*⟩_box_ and 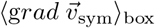. We also simulate the possible evolutions of ⟨*ε_cell_*⟩_box_ by injecting in Eq. 1 the experimental value of *grad* 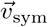 and by varying the parameter *τ_r_* (Fig. 11).

In Fig. 4c, we compare the experimental measurement of deformation evolution, ⟨*ε_cell_*⟩_box_ (red curve), with various simulations: we vary *τ_r_* between 10 minutes (light grey) and infinity (black), the latter being a purely elastic limit (Fig. 11). The simulated deformation *amplitude* agrees reasonably with the experiment when we use *τ_r_* ≈ 20 min. The Fourier determination of deformation is sufficiently discriminant and reproducible to determine correctly the amplitude order of magnitude and we can reasonably exclude values of τ_r_ outside of the interval [15 min, 25 min] (Fig. 12).

Thus measurements coarse-grained at the cell group scale yield an estimate of an effective cell deformation relaxation time. Since divisions play a negligible role on the experiment timescale, the effective cell deformation relaxation time is necessarily associated with dynamics of cell rearrangements, which are numerous in particular near the constriction entrance (Figs. 4d, 13, Supp. Movie 5)

This relaxation time value is of same order of magnitude than the one we found for MDCK cells in a 2D migration experiment of cells around an obstacle: *τ* ≈ 70 min (30). Assuming that the effective viscosity at cell group scale due to rearrangements is

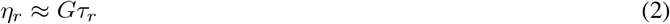

and taking *η_r_* ≈ 10^5^ Pa.s as we found previously (6, 9), Eq. 2 yields *G ≈* 100 Pa. Since *G* is not expected to vary with scale, so is the same at cell scale and at cell group scale. It is also expected to be close to the modulus of an isolated cell outside of an aggregate. In fact, the isolated F9 cell modulus is 100 Pa too (31).

Note that the determination of *τ_r_* in Fig. 4c is not sensitive to the *shape* of the deformation evolution. The peaks in deformation are not systematically observed and their origin is unknown; they are likely due to a stick-slip friction on the glass coverslip (rather than to cascades of rearrangements, which we do not detect). Determining accurately the shape of the deformation evolution is beyond the scope of the present paper. It would require to improve simultaneously the time resolution and signal-to-noise ratio beyond the possibilities of the current Fourier method with our current image quality. This would be possible in principle using cell contour segmentation complemented with detailed tensorial analysis.

### 3.3 Cell rearrangements dynamics

Using the same data, but at cell scale, we have access to the cell rearrangements dynamics at the scale of a single event (Fig. 5a) by manually identifying and segmenting cell-cell contacts which are disappearing and newly created.

**Figure 5:**
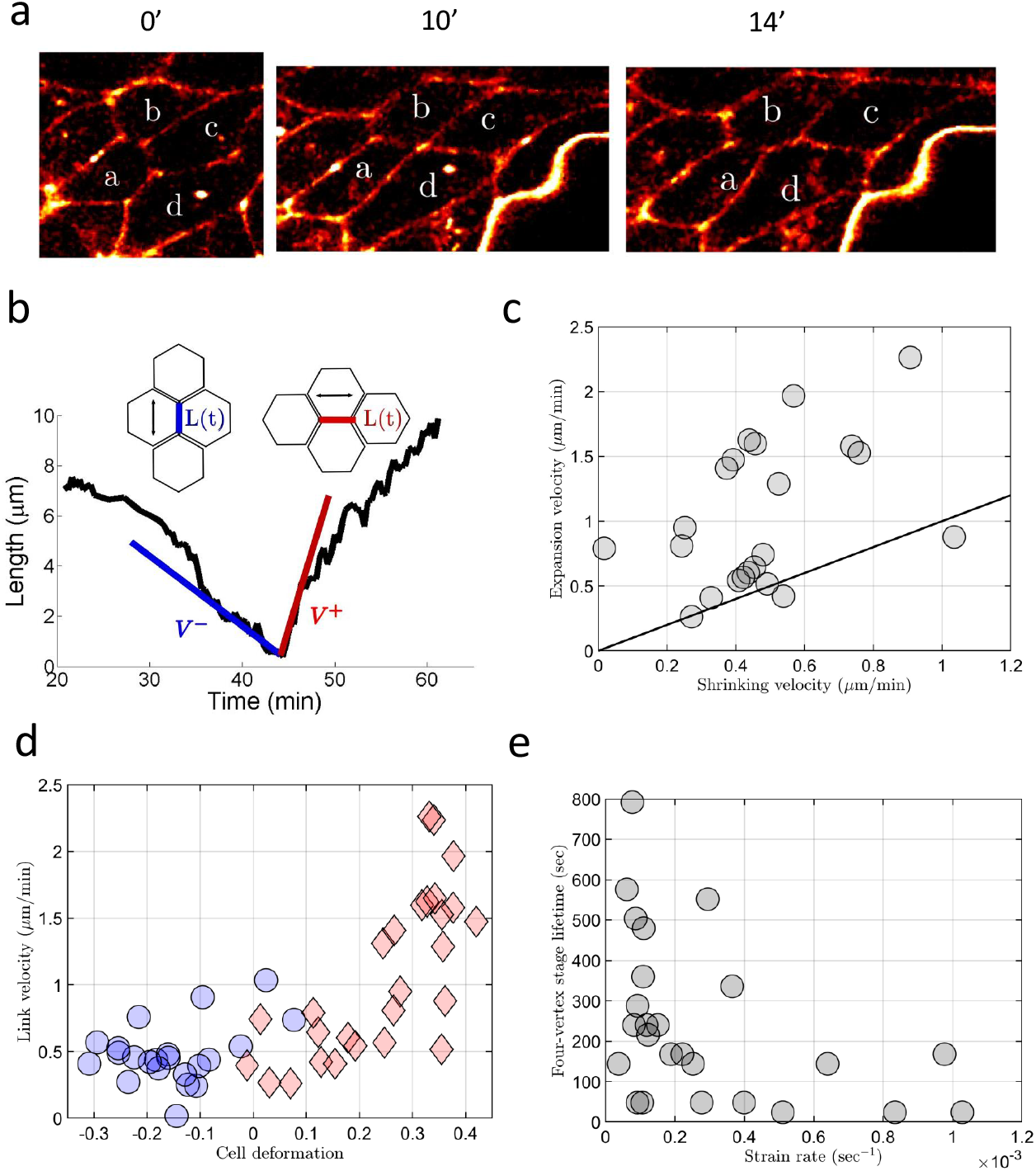
Cell rearrangements. (a) Typical cell rearrangement in the aspired aggregate. Before the rearrangement, cells *a* and *c* are in contact, this contact progressively disappeared until reaching the four-fold vertex stage. After the rearrangement, a new contact between cells *b* and d is created. Times are indicated in minutes. (b) Junction length evolution during a rearrangement. Length of the disappearing junction (in blue) and of the newly created one (in red) versus time, see inset for definitions. (c) Asymmetry in junction length evolution speed before and after the four-fold vertex: velocities of creation versus disappearance. (d) Link shrinkage and growth versus deformation: *V*^-^ versus 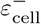 (blue) and *V*^+^ versus 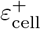 (red); see text for definitions. (e) Stability of four-fold vertex: time *T*_vertex_ where the junction that undergoes a rearrangement is below a cutoff length *L*_small_ = 1.5 *μ*m, versus 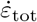.

As the deformation rate and cell deformation fields are heterogeneous, we can trigger in the same experiment: (i) “quasi-static” rearrangements in deformed regions of the tissue away from the channel, where both a disappearing link between cells losing contact and a newly appearing link between cells gaining a new contact can be clearly identified; and (ii) “sliding” rearrangements at the vicinity of the channel, where cells are highly stretched, flow rapidly and seem to slide on each other. In the first case where rearrangements can be clearly identified and well-defined, we manually measured the disappearing and appearing junction lengths versus time (Fig. 5b). We define velocities of appearance and disappearance before and after the rearrangement, by taking the tangent of the junction lengths curves before and after the four-fold vertex stage. To have a robust definition, we define the velocity of disappearance *V*^-^ = *L*^-^(*t* = -*T*)/*T* and of creation *V*^+^ = *L*^+^(*t* = +*T*)/*T* with *T* = 6 min. Their values are unchanged if we take *T* ~ 10 min, as the length evolution is locally linear around the four-fold vertex stage. We often see an asymmetry in *V*^+^ and *V*^-^ (Fig 5c).

To relate these quantities with the local cell deformation during the rearrangement, we project the tensor *ε_cell_* on the direction of the junction that disappears / appears. It yields 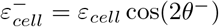 where *θ*_-_ is the angle between the disappearing link and the principal direction (i.e. eigenvector corresponding to the positive eigenvalue) of *ε_cell_*; and 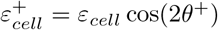 where *θ*^+^ is the angle between the appearing link and the principal direction of *ε*_cell_ (Fig 5d). We observe that the disappearing junction is compressed, and that the newly created junction is elongated, which is consistent with the fact that observed rearrangements are triggered by the cell group scale flow imposed by the aspiration.

We plot *V*^+^ and *V*^-^ versus 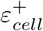 and 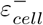, that are averaged on the same time *T* = 6 min as the velocities in Fig. 5d. For −0.3 < *ε_cell_* < 0.3, the velocities *V*^-^ and *V*^+^ are around the mean value ≈ 0.5 *μ*m/min. For 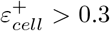, the mean value of *V*^+^ increases suddenly up to ≈ 2.5 *μ*m/min (see also Supp. Movie 5). Examples of such highly deformed rearranging cells are visible in Figs. 4d and 5a; they occur mainly near the constriction entrance. They are usually compatible with the tissue confluence and integrity. High aspiration pressures and velocity can result in aggregate fractures (Supp. Movie 6): we do not analyse these experiments.

We also looked at the life time T_vertex_ of a “four-fold vertex stage” (Fig. 5e). We define it as the stage where the cell-cell junction is below 1.5 *μ*m, which is small compared to the cell size, and large enough to avoid artefacts due to the resolution on the length determination when the junction is small. We observe that the life time of a four-fold vertex at null deformation rate can be large, which is qualitatively different from the physics of liquid foams, where the four-fold vertex is unstable (32). The life-time of the four-fold vertex in a biological tissue could be related to the dynamics of adhesion remodeling (33, 34), i.e. the timescale necessary for non-contacting cells to create a new junction. At a high cell deformation but null deformation rate, we could imagine that the four-fold vertex lifetime is dominated by active fluctuations and adhesion remodeling. Conversely, at both high cell deformation and deformation rate, a new contact between cells can be created without maturing cell-cell adhesion (i.e. new contact and sliding of cells are triggered by the deformation rate).

### 3.4 Aggregate relaxation after aspiration

The aspired aggregate can either relax in a reservoir at the end of the channel or being pushed back in the entrance reservoir. In both cases, the relaxation remains an actual two-dimensional relaxation as the tissue is still confined in the z direction. The relaxation dynamics qualitatively depends on the time the tissue has spent in the channel, and two regimes can be distinguished.

During a rapid aspiration, cell rearrangements do not have time to occur and the aggregate deformation is uniquely due to cell deformation. In this case, the aggregate shape relaxation in a reservoir corresponds to a cell deformation relaxation on the minute timescale (Fig. 6, Supp. Movie 7). This corresponds to an effective visco-elastic solid behavior (Kelvin-Voigt) with a typical timescale of relaxation

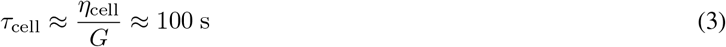

**Figure 6:**
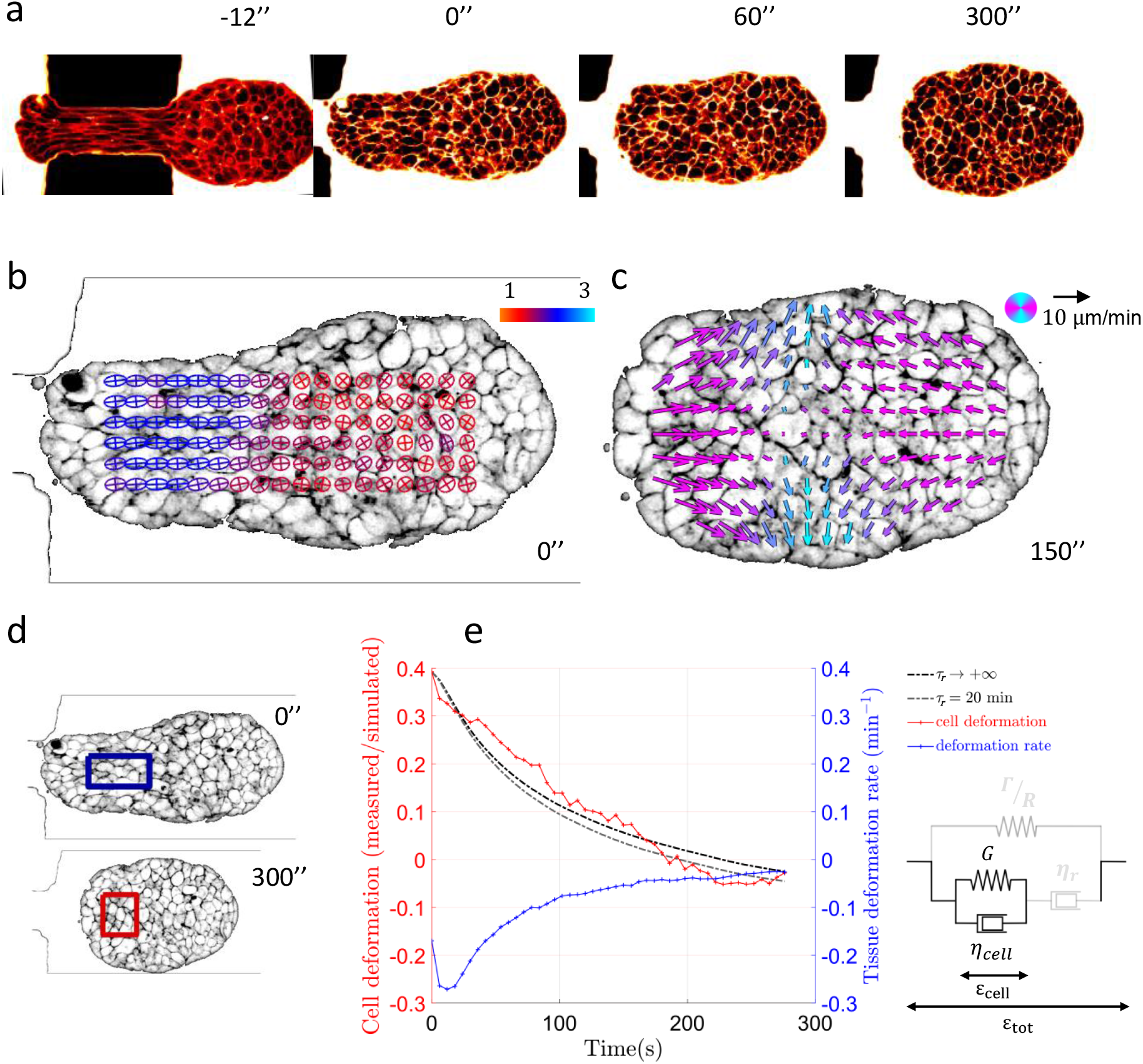
Aggregate rapid relaxation. (same analysis and imaging methods as in Fig. 4). (a) Typical aggregate during rapid relaxation phase (Supp. Movie 7). The aggregate is aspired in a short channel, the right part had time to slowly relax during the aspiration. When the radius of the remaining part of the aggregate to be aspired is of the same size than the channel size *(t* = −12*s*), the aggregate part still aspired within the channel relaxes quickly *(t* = 0*s*) until the aggregate reaches a symmetric shape (*t* ≈ 200s). (b) Ellipses representing cell group anisotropy, color-coded from 1 (red) to 3 (blue). (c) Velocity field during the relaxation. (d) Virtual patch of tissue advected and deformed by the velocity field and its gradient, measured by optic flow. (e) In blue, anisotropic component of cell group deformation rate 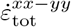, in red 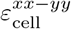, in gray levels simulated 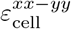 for different typical values of cell shape relaxation timescale *τ_r_*, in black simulated 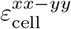 if there was no cell shape relaxation (purely elastic deformation, *τ_r_* → ∞). Times are indicated in seconds.

Taking *G* ~ 100 Pa, Eq. 3 yields: η_cell_ ~ 10^4^ Pa.s.

If the aspired aggregate is aspired quasi-statically at a very slow pace (*T*_aspi_ > *τ_r_*, as is the case in Fig. 4), the cell deformation has enough time to relax through rearrangements (Fig. 7). In this case, when the aggregate starts to relax in the reservoir, the cells shape is already isotropic but we still observe a relaxation of the aggregate shape (Fig. 7a, Supp. Movie 8) associated with an elongation of cells along the axis perpendicular to the aggregate main elongation axis. This phenomenon can be explained by the fact that aggregate surface tension triggers a compaction of the aggregate along its main axis of elongation. Since cells can rearrange and relax their shape under stress on timescales *τ_r_* ~ 20 minutes, the aggregate surface tension related compression triggers significant cell deformations without any rearrangement on timescales shorter than *τ_r_*. This is the signature of an elasto-capillar behavior for the tissue, which can happen when the aggregate surface tension Γ is high enough to trigger significant cell deformations. We assume here that the aspiration does not modify the order of magnitude of Γ, which is Γ = 5 mN/m according to our previous aggregate scale experiments using F9 cells (35). The typical aggregate curvature radius R at the end of the rounding phase is of order of R_channel_/2 ~ 50 *μ*m. We obtain:

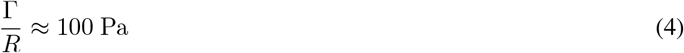

**Figure 7:**
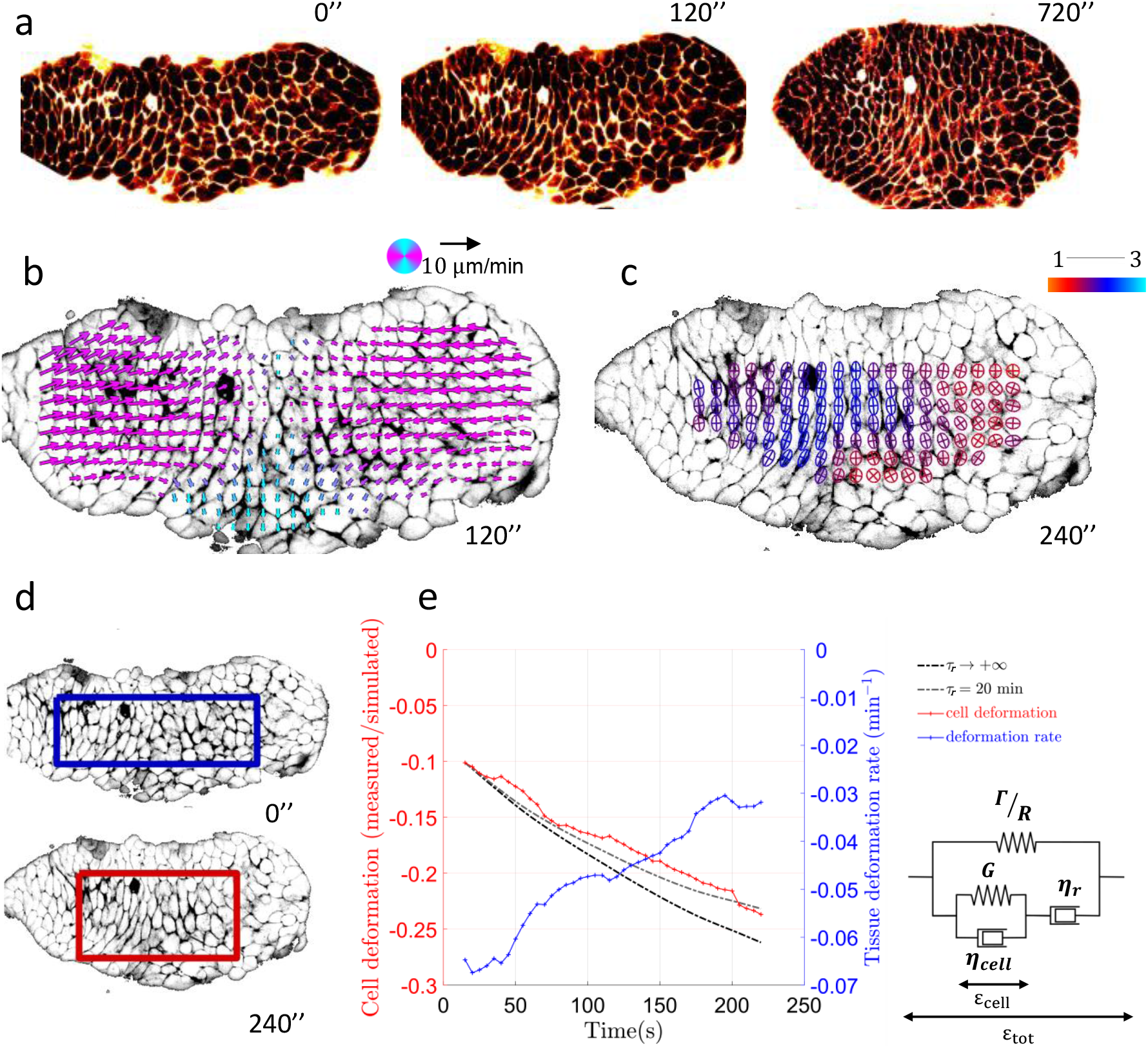
Aggregate long-term relaxation. same analysis and imaging methods as in Fig. 4). (a) Typical aggregate during long-term relaxation phase (Supp. Movie 8). The aggregate is aspired in a long channel. When the aggregate is released from the channel after a longer time spent in the channel (30 minutes), it undergoes first a rapid and partial shape relaxation similar to the one of Fig. 6, associated with cell elongation relaxation along the *x* axis, and then continues to relax by compressing the cells along the x axis. (b) Velocity field corresponding to the compaction of the aggregate obtained with optic flow. (c) Ellipses representing cell group shape anisotropy coded from 1, red, to 3, blue. (d) Virtual patch of tissue advected and deformed by the velocity field and its gradient, measured by optic flow. (e) In blue, anisotropic component of cell group deformation rate 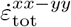, in red 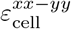, in gray levels simulated 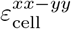 for different typical values of cell shape relaxation timescale *τ_r_*, in black simulated 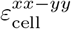 if there was no cell shape relaxation (purely elastic deformation, *τ_r_* → ∞). Times are indicated in seconds.

This value is of same order of magnitude than the elastic modulus G, which confirms that the capillarity is sufficiently large to induce high elastic deformations within the bulk of the aggregate. Fig. 14 shows an example of aggregate relaxing after being partially aspired sufficiently slowly in a longer channel (Supp. Movie 9). After the fast cell shape relaxation, the aggregate does not entirely come back yet to its initial round shape. The aggregate then rounds up due to capillarity, while cells re-deform, within a few minutes (Supp. Movies 10, 11). Myosin is essential in this shape relaxation process since when blebbistatin is added, the relaxation is partial: the first fast relaxation is conserved, but not the second slow one (Fig. 15, Supp. Movie 12).

In turn, this new elastic deformation can eventually fully relax. This occurs at low shear rate, and long timescales, where cell divisions may also play a role.

To recapitulate the different behaviors observed in the previous sections, we propose a multi-scale rheological diagram of the aggregate (Fig. 8 and Table 1). It combines the contributions of intra-cellular rheology (cell deformations), inter-cellular rheology (cell rearrangements) and aggregate surface tension (elasto-capillarity). This complete rheological model could in principle apply to all experiments presented here (Figs. 4c, 6e, 7e). Ideally, all of them could be fitted with a single set of parameters, at least if elastic and viscous moduli are supposed constant. We however emphasize that only the aspiration part of the experiment is here submitted to a quantitative test. The relaxation part could be tested more quantitatively through a finite element simulation taking into account the geometry details.

**Figure 8:**
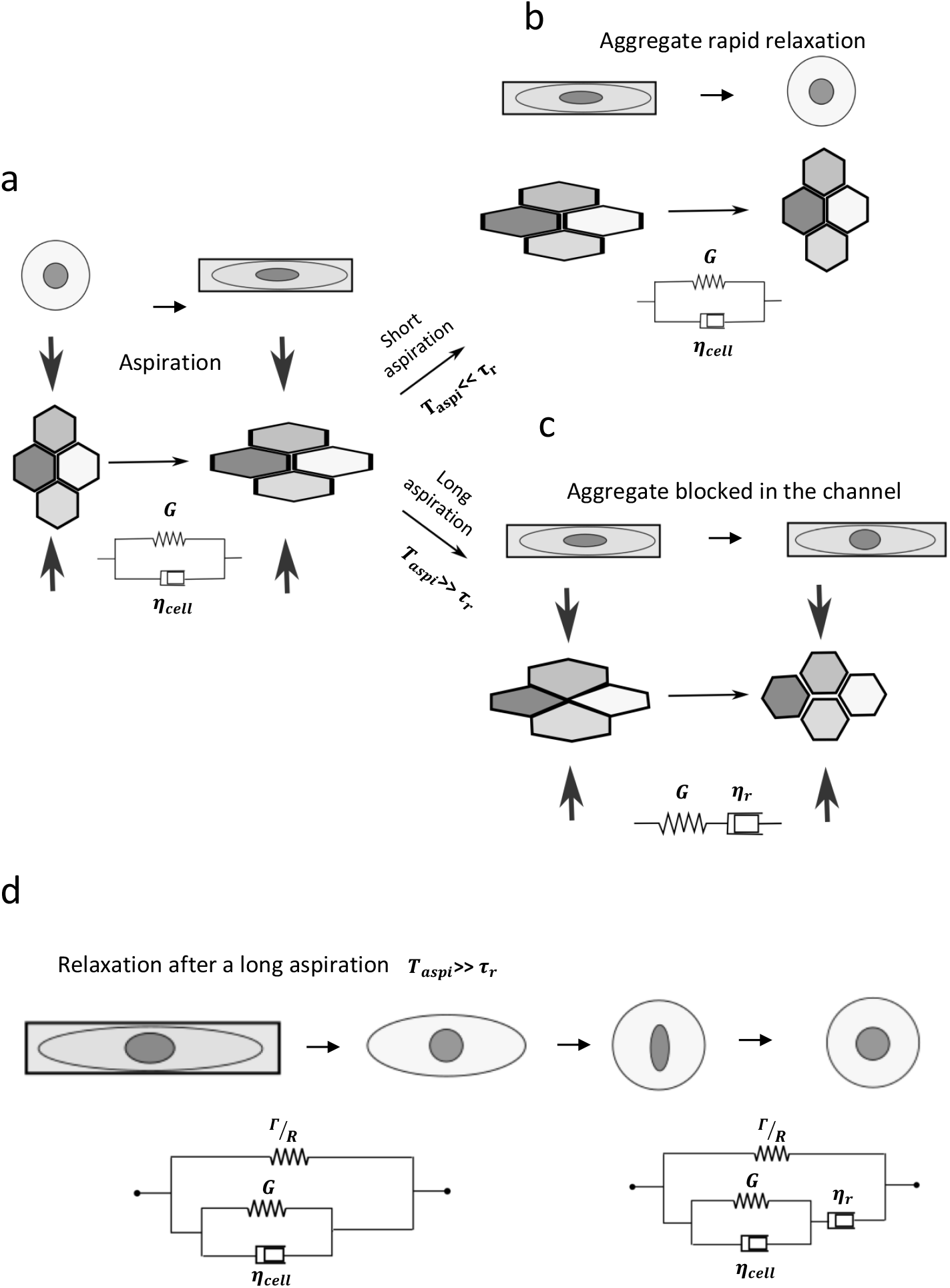
Proposed rheological diagram for the aggregate. (a) Cell group without rearrangements: cells can be described as a visco-elastic solid (Kelvin-Voigt) with a typical timescale of cell shape relaxation *τ_cell_*. (b,c) Schematic representation of what happens during an aspiration and a relaxation experiments depending whether the cell shape has time to relax due to cell rearrangements during the aspiration phase: *T_aspi_* either smaller (b) or larger (c) than the relaxation time τ_r_. In (c), the tissue can be at first order described as a visco-elastic liquid (Maxwell) with a relaxation time *τ_r_*. (d) Schematic rheological diagram proposed for the whole aggregate, with Γ/*R* an aggregate scale effective capillary modulus as the surface tension drives the aggregate’s rounding, until eventually the whole elastic deformation fully relaxes at long timescales. See Table 1 for parameter names and orders of magnitude.

## 4 Conclusion and perspectives

### 4.1 Summary

Our experimental set-up uses a microfluidic constriction to impose a highly heterogeneous flow to a 3D cell aggregate, by controlling both the force applied to the aggregate and the flow geometry. It triggers (i) cell deformations, (ii) cell rearrangements and (iii) global aggregate geometry changes, and enables to study the interplay between the three. Using two-photon microscopy we simultaneously image cell contours at sub-cellular resolution and quantify mechanical response at the scales of cell, cell group and whole aggregate.

While the basic principle is similar to micropipette aggregate aspiration, the present set-up has at least three advantages. First, we can study separately, and in detail, the effect of cell group elasticity, aggregate surface tension, and coupling between both (so-called “elasto-capillarity”). Second, we can reach higher cell deformation, or more precisely we obtain a large product of deformation rate by relaxation time (so-called “Weissenberg number”, here of order of one); this is probably relevant

**Table 1:** Characteristic quantities for the proposed rheological model, and their approximate order of magnitude (Fig. 8).

| *quantity* | *symbol* | *value* | *source* |
| --- | --- | --- | --- |
| cell group scale visco-elastic relaxation time | $\tau_r$ | $10^3$ s | Fig. 4c |
| cell group scale viscosity | $\eta_r$ | $10^5$ Pa.s | Refs. (6, 9) |
| elastic modulus | $G$ | $10^2$ Pa | Eq. 2 |
| cell scale visco-elastic relaxation time | $\tau_{\text{cell}}$ | $10^2$ s | Figs. 6e, 7e |
| cell scale viscosity | $\eta_{\text{cell}}$ | $10^4$ Pa.s | Eq. 3 |
| aggregate scale capillary modulus | $(\Gamma/R)$ | $10^2$ Pa | Eq. 4 |

in fast morphogenetic changes phases like gastrulation, where tissues encounter both high cell deformations and cell deformation rates. Third, we can map different scales and local differences in mechanical properties; we are thus not restricted to homogenenous aggregates.

### 4.2 Perspectives for biophysics

3D cell aggregates are model systems of active fluctuating cellular materials. We propose here to measure tissue effective rheological parameters averaged on time and space. Improvements could include changes in constriction walls, which could be rounded or at angles smaller than 90 degrees. Embedding deformable stress sensors inside the tissue (20, 36–38) would enable to relate cell deformation with the local tissue stress to validate further rheological models; beyond the linear models we test here, future works might investigate more realistic power-law rheologies.

Further progresses could come from improved time and space resolutions, segmentation and tracking, to obtain better statistics; dynamic stress inference (which applies even to moving junctions) (39, 40) could infer junctions tensions and determine how they are modulated by cell deformations and rearrangements.

How cell membrane trafficking, adhesion proteins and molecular motors dynamics affect rearrangements dynamics and are relocalized during junction remodeling in 3D tissues is still not well understood (41, 42). Combining our method with cell lines endowed with fluorescent reporters of such proteins would enable to quantify their dynamics using fluorescence recovery after photobleaching technics for example. Other exciting insights could come from blocking the action of some proteins (using knock-out cell lines or chemical inhibition of various pathways and effector proteins), so as to quantify the relative contributions of various components of the cellular machinery to tissue mechanics. Studying the effects of drugs such as blebbistatin, which inhibits Myosin II could also brings new interesting pieces of information, as shown by our proof of concepts experiments, see Fig. 15b, Supp. Movie 12. In the same way, cell-cell stress propagation can also be modified for instance using α-catenin null cell line. Since α-catenin links the cytoskeleton with cadherins, cell-cell junctions are then modified. Aggregates still form cadherin-cadherin junctions, but cells shape are more irregular and there is a shorter correlation range in both shape and velocity, while rearrangements are dramatically impeded (Supp. Movie 13).

### 4.3 Perspectives for developmental biology, spheroids and organoids

Cell rearrangement dynamics has been studied in *in vivo* systems like Drosophila (15, 16, 33, 34, 43) or Xenopus embryos (44), *in vitro* tissues (30) and numerically for *in silico* tissues (45). However, in the context of embryo development or collective cell migration *in vitro*, it is difficult to distinguish the rearrangements which build or relax deformation (or stress) (34), i.e. those where cells act as motors, versus those where cells resist a force imposed from outside. Here, we present an experimental platform suitable to quantitatively characterize the resistant cell rearrangements triggered by an imposed flow.

This microfluidic platform could also be a useful simple tool for perturbing mechanosensitive pathways by applying shear stresses and to quantify their effect on biomechanical networks (using fluorescent reporters or doing RNA sequencing afterwards, as it is easy to collect the aspired aggregates after the experiment). For this kind of applications, it could be useful to develop parallel and/or sequential constriction set-ups, as has already been done for single cells experiments (46).

It has been recently shown that gradients of expression of cadherins are associated to embryonic organoids symmetry breaking and elongation (47). Whether these gradients are associated to gradients in rheological properties remains unknown in such systems. An advantage of our method is that it relies on the observation and extraction of local mechanical response in 3D: it could be used to identify spatial heterogeneities in cell group mechanical response in systems such as embryonic organoids and correlate them with local protein or gene expression.

This setup will be very useful for understanding the mechanisms at stake in the response of 3D tissues to mechanical stresses and in particular to get new insights in the biophysics of cell-cell rearrangements which plays a central role in development. We envision that it could be used in many different systems such as Hydra (from which aggregates can easily be formed), mouse embryos, organoids, 3D bulk tissues of Xenopus, and could yield new information on key developmental processes.

## Acknowledgements

We thank the Nanoptec platform at ILM and especially Christophe Moulin, and the ImagoSeine core facility. We thank A. Nagafuchi for his generous gift of F9 cells, David Charalampous for developing the pressure control device, Alain Richert for help on tissue culture, Rémy Fulcrand for help on wafer fabrication in clean room, Pascal Hersen for letting us use his wet lab for microfluidic device preparation.

All home-made codes and datasets are available upon reasonable request to the corresponding author.

## Supplementary Material

### Supplementary Figures

**Figure 9:**
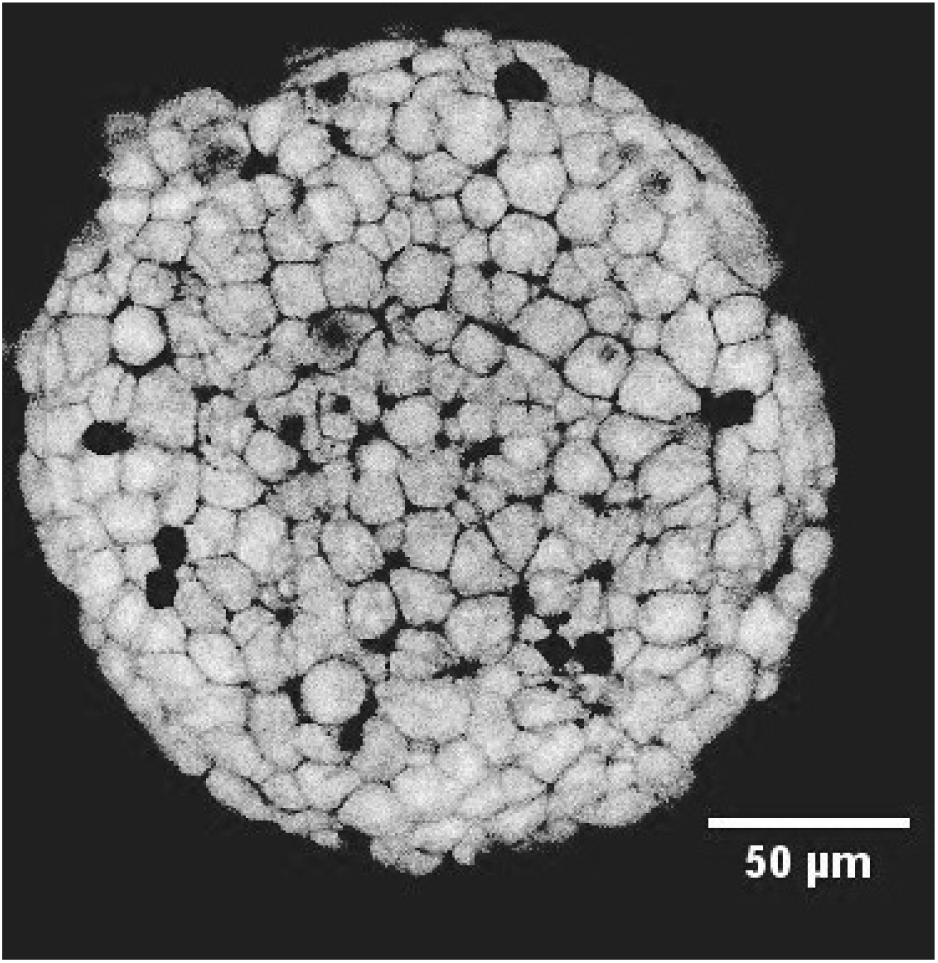
F9 cell aggregate mid-section (two-photon microscopy).

**Figure 10:**
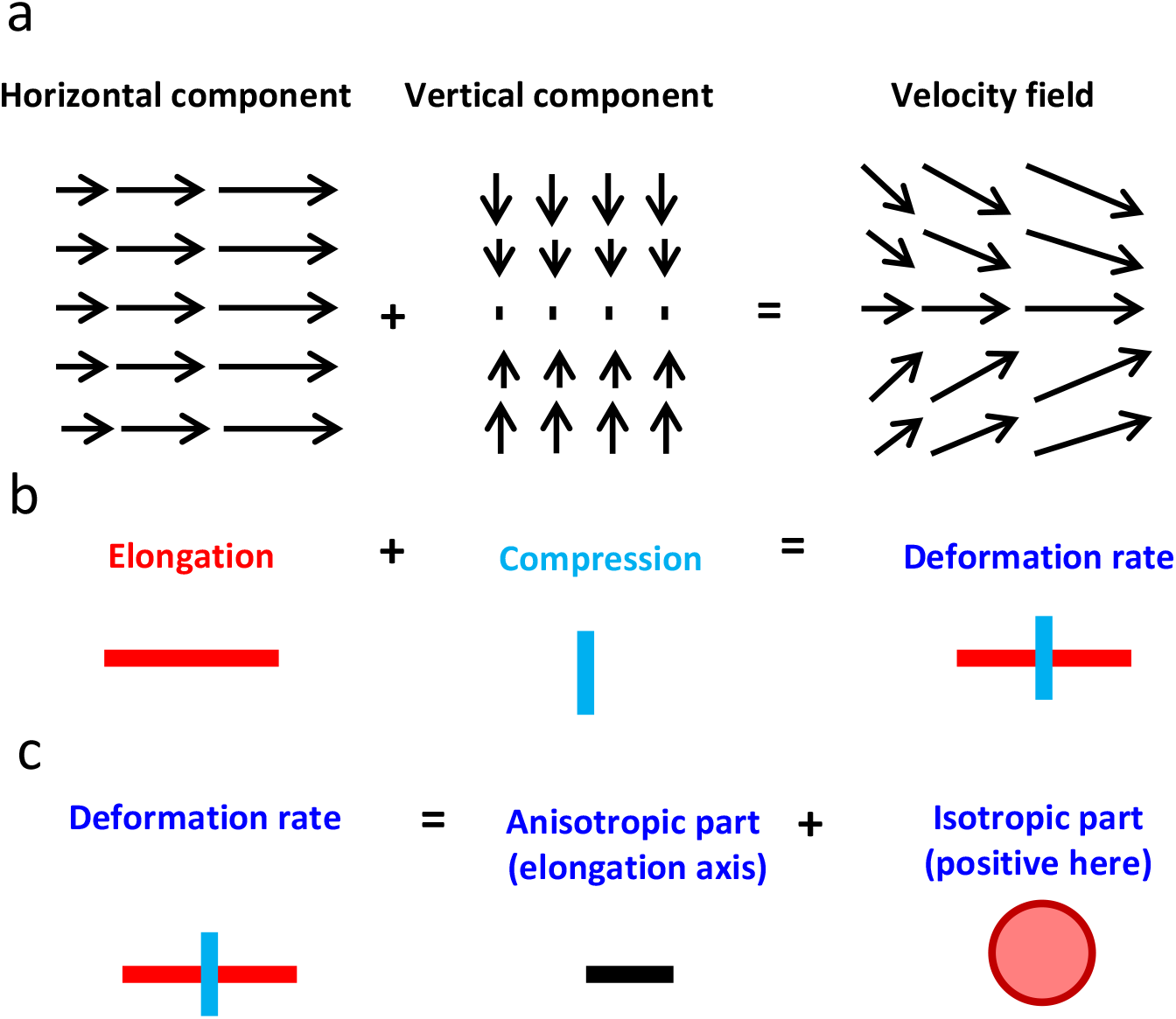
(a) Decomposition in horizontal and vertical components of a typical velocity field in the constriction. (b) Corresponding graphical representation of deformation rate (15). (c) Decomposition of the deformation rate in anisotropic and isotropic parts, and corresponding graphical representation (15).

**Figure 11:**
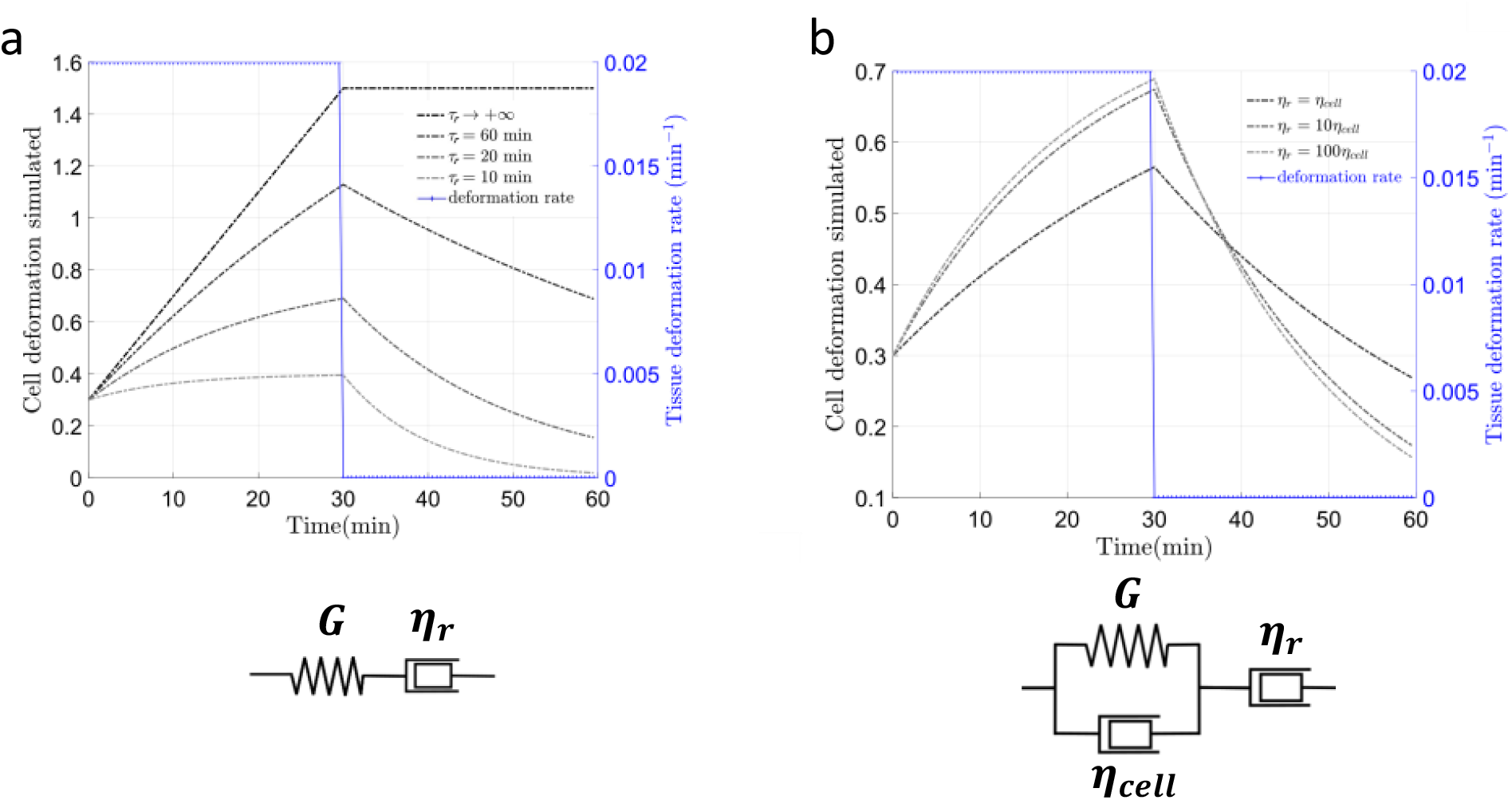
Idealized mechanical responses. Simulated time evolutions for different models (insets in a,b), *τ_r_* values and viscosity ratios of the response to an imposed deformation rate step. Simulated equations are for (a) 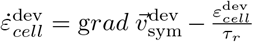 and for (b) 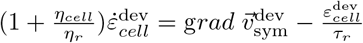.

**Figure 12:**
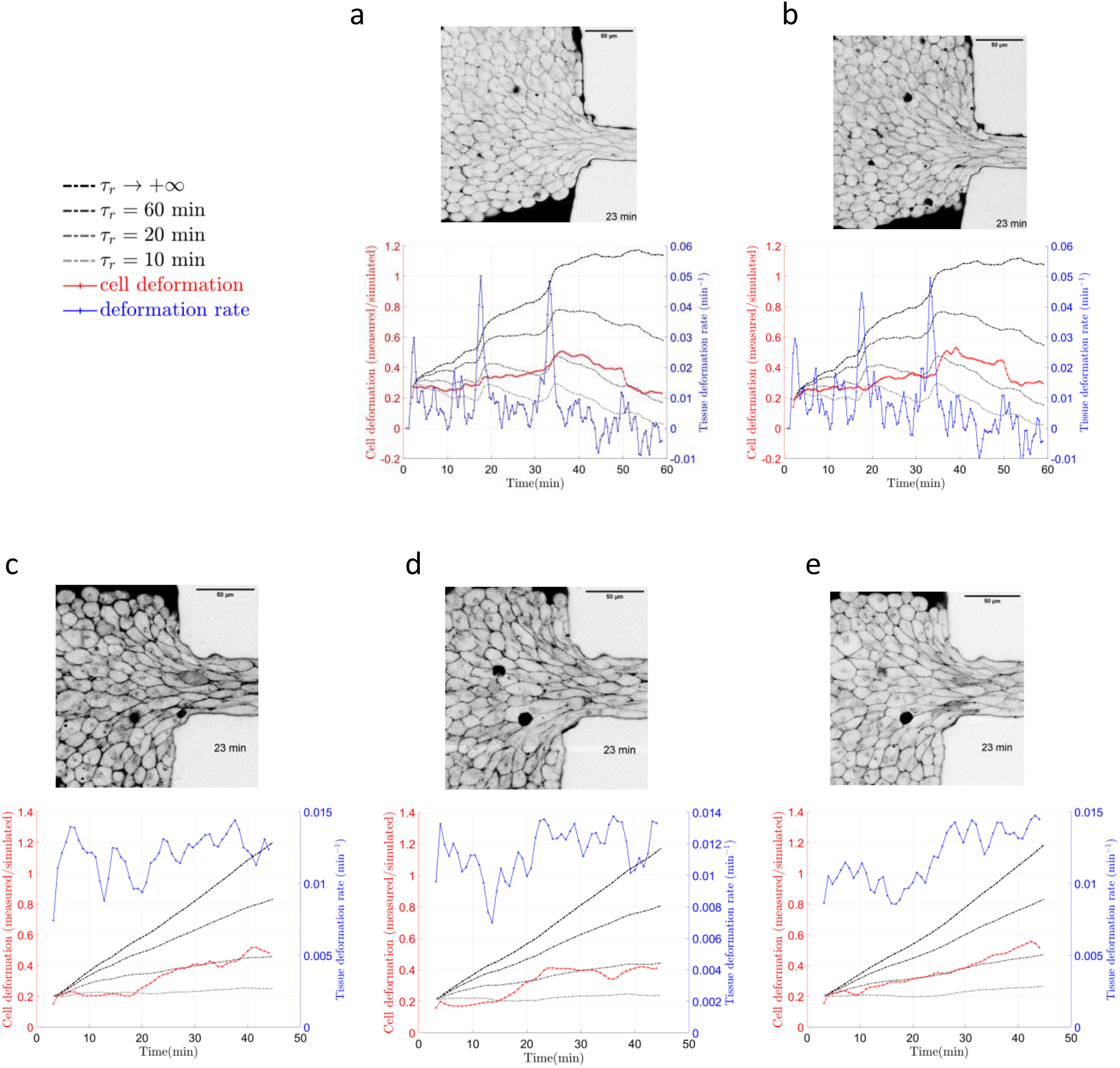
Reproducibility in estimating. *τ_r_* (same analysis and imaging methods as in Fig. 4). (a,b) Same experiment as in Fig. 4, imaged and analyzed at two different heights (Supp. Movie 14); (a) is Fig. 4c duplicated here to enable comparison taken at *z* = 15 *μ*m from the coverslip and (b) at *z* = 30 *μ*m from the coverslip. (c-e) Another experiment, imaged and analyzed at three different heights (Supp. Movie 15) (Heights: (c) *z* = 20 *μ*m, (d) *z* = 30 *μ*m and (e) *z* = 40 *μ*m from the coverslip).

**Figure 13:**
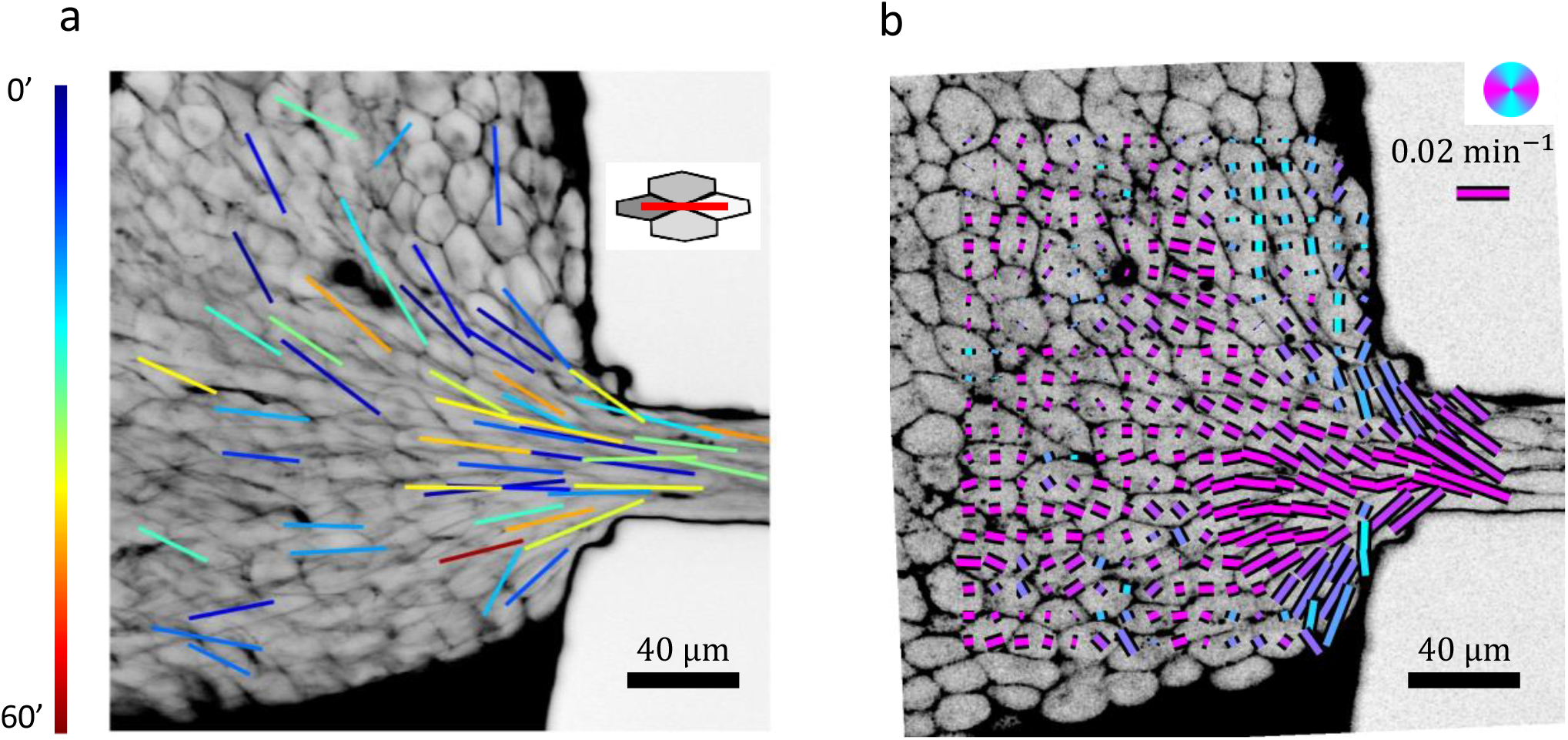
Cell rearrangement map. Rearrangements are detected in Supp. Movie 4 during one hour. (a) Position and orientation of the rearrangements detected, plotted as a bar linking both centers of cells that will lose contact just before the rearrangement (inset), and color-coded according to the timing of the four-vertex stage (in minutes). (b) Deformation rate anisotropic part averaged over time (60 minutes) with the bars color coding for their angle with respect to the channel. Zones of high rearrangements frequency correspond to high deformation rate zones.

**Figure 14:**
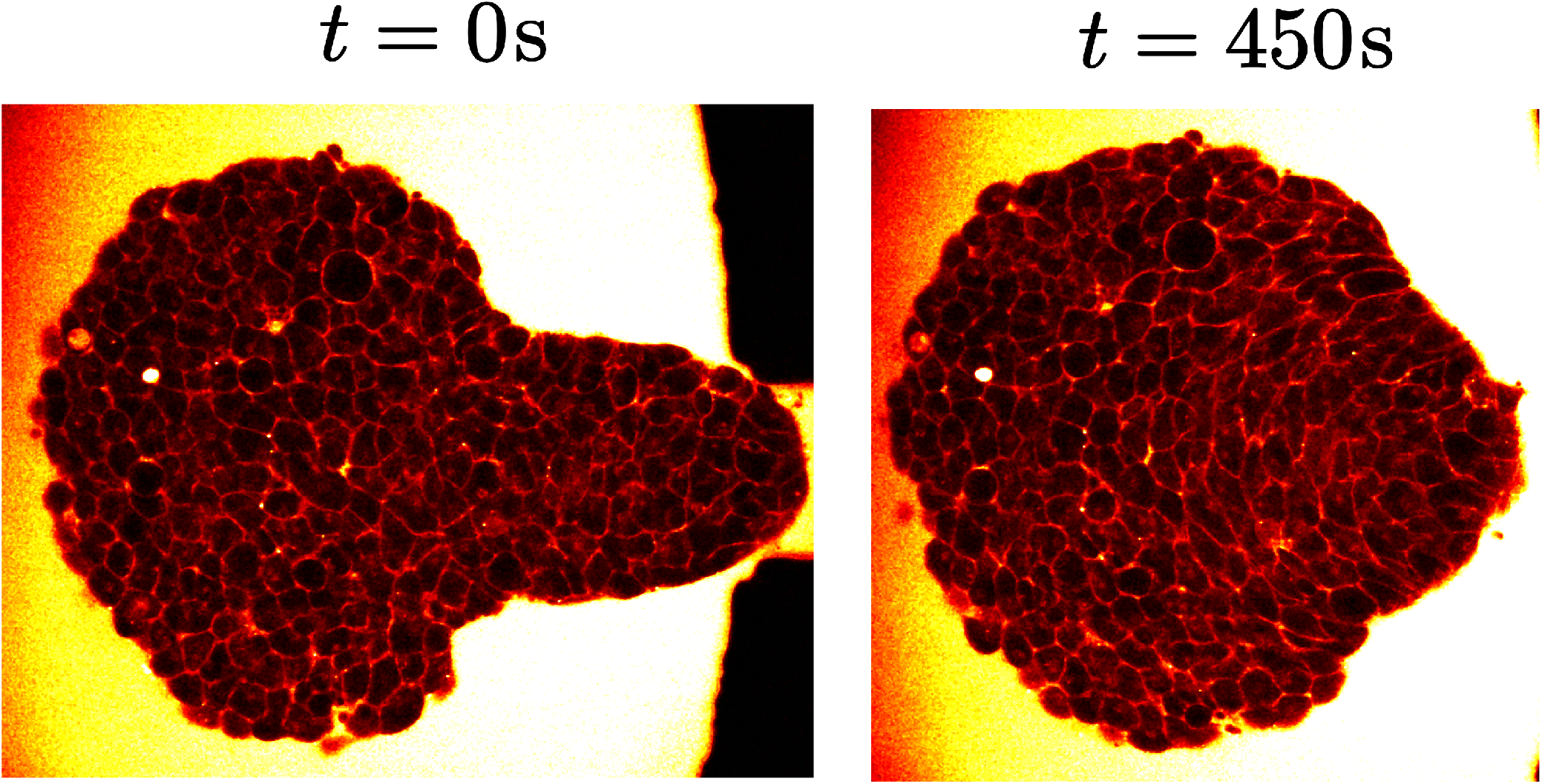
Aggregate compaction after a relaxation. An aggregate has been aspired during 1000 s. It is pushed back in the reservoir and cell shapes relax within a few seconds. Time *t* = 0 corresponds to 20 s after the pushing and cell shapes have already almost totally relaxed. After 450 s, the aggregate as a whole is round, but not each cell individually (Supp. Movie 9).

**Figure 15:**
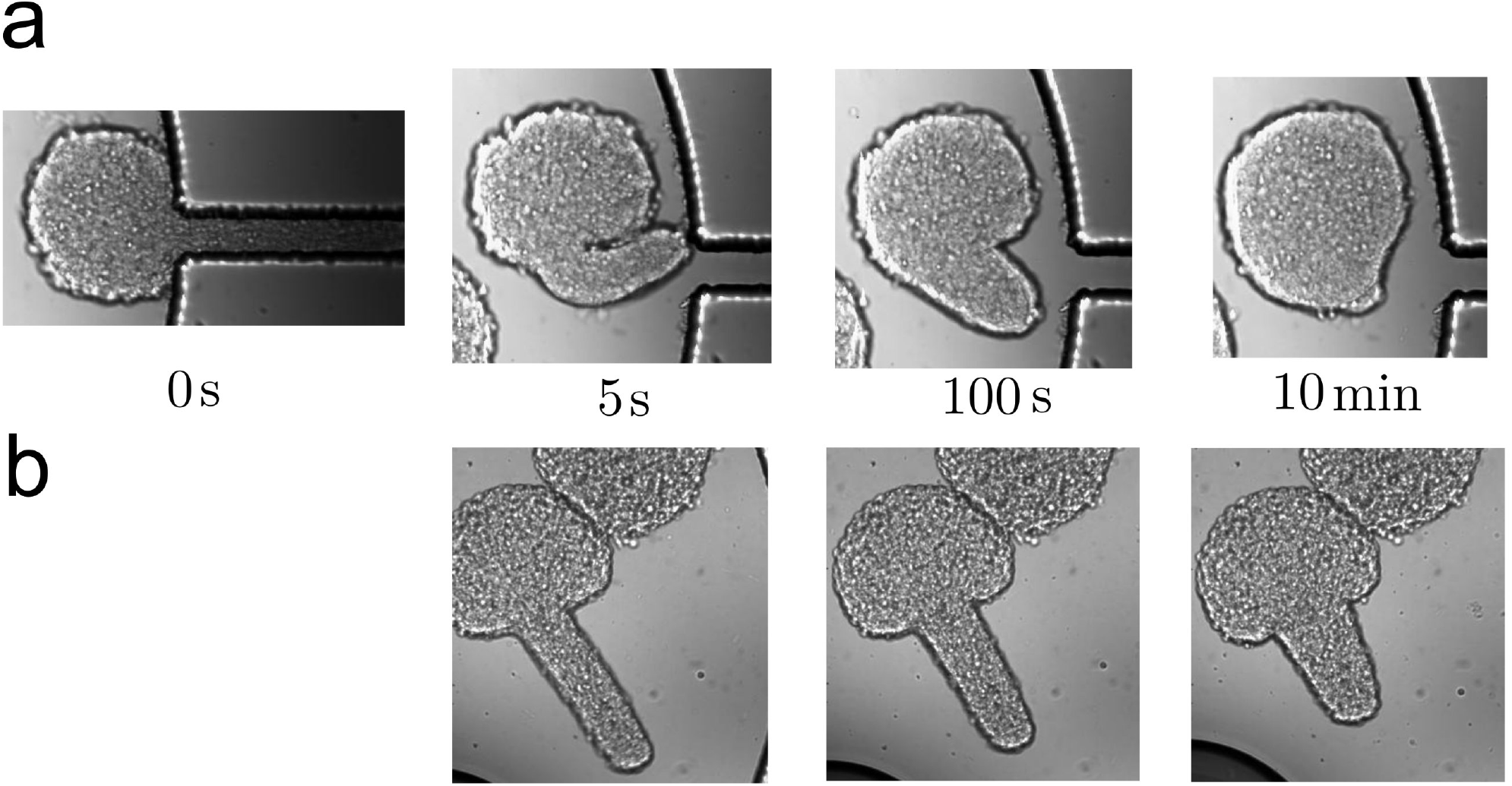
First and second phases of relaxation. (a) Brightfield image of aggregate relaxation. After a 1000 seconds long aspiration, the aggregate is left free to relax. Time *t* = 0 is counted at the end of the aspiration, *i.e*. at the beginning of the relaxation. (b) Same with 50 **μ**M of blebbistatin to inhibit myosin II. The first fast phase of the relaxation is still present, but there is no complete return to the initial shape.

### Supplementary Movies

Supplementary movies can be downloaded from the following link: https://doi.org/10.5281/zenodo.6895913

Supp. Movie 1: 40 minutes long timelapse (brightfield) of an aggregate aspired in a channel.

Supp. Movie 2: z-stack (two-photon microscopy) of a non-deformed F9 cells aggregate.

Supp. Movie 3: 60 minutes long timelapse (two-photon microscopy) of an aggregate aspired in a channel. Left: imaged plane at *z* = 40 *μ*m from the coverslip. Middle: imaged plane in contact with the coverslip. Right: superposition of the two imaged planes.

Supp. Movie 4: 60 minutes long timelapse (two-photon microscopy) of an aggregate aspired in a channel.

Supp. Movie 5: cell-cell junctions involved in cell rearrangements in Supplementary Movie 4. Rapid relaxation rearrangements (*V*^+^ > 1 *μ*m/min) are represented in red while slow ones (*V*^+^ < 1 *μ*m/min) are in black.

Supp. Movie 6: tissue fracture appearing during a rapid aspiration (two-photon microscopy timelapse).

Supp. Movie 7: aspiration and rapid relaxation experiment (two-photon microscopy).

Supp. Movie 8: relaxation experiment after an aggregate being fully blocked in the aspiration channel during 30 minutes (two-photon microscopy).

Supp. Movie 9: relaxation experiment after a partial aspiration during which cellular deformation has relaxed (two-photon microscopy).

Supp. Movie 10: relaxation experiment after an aspiration in a short channel, with a more complex geometry during the rounding phase (two-photon microscopy).

Supp. Movie 11: other example of a relaxation experiment after an aspiration in a short channel (two-photon microscopy).

Supp. Movie 12: relaxation experiment after a partial aspiration (two-photon microscopy) in 50 *μ*M blebbistatin medium.

Supp. Movie 13: aspiration of an α-catenin null cell line aggregate (two-photon microscopy).

Supp. Movie 14: same aspiration as in Supp. Movie 4, with two planes superimposed in red and green (heights: *z* = 15 and 30 *μ*m from the coverslip).

Supp. Movie 15: aspiration with three planes superimposed in red, blue and green (heights: *z* = 20, 30 and 40 *μ*m from the coverslip).

